# Inferring single-cell spatial gene expression with tissue morphology via explainable deep learning

**DOI:** 10.1101/2024.06.12.598686

**Authors:** Yue Zhao, Elaheh Alizadeh, Hash Brown Taha, Yang Liu, Ming Xu, J. Matthew Mahoney, Sheng Li

**Affiliations:** The Jackson Laboratory for Genomic Medicine, Farmington, CT, USA; Department of Cancer Biology, Norris Comprehensive Cancer Center, Keck School of Medicine, University of Southern California, Los Angeles, CA, USA; Institute on the Biology of Aging and Metabolism, University of Minnesota, Minneapolis, MN, USA; Department of Biochemistry, Molecular Biology & Biophysics, University of Minnesota, Minneapolis, MN, USA; The Jackson Laboratory for Mammalian Genetics, Bar Harbor, ME, USA

## Abstract

Deep learning models trained with spatial omics data uncover complex patterns and relationships among cells, genes, and proteins in a high-dimensional space. State-of-the-art *in silico* spatial multi-cell gene expression methods using histological images of tissue stained with hematoxylin and eosin (H&E) allow us to characterize cellular heterogeneity. We developed a vision transformer (ViT) framework to map histological signatures to spatial single-cell transcriptomic signatures, named SPiRiT. SPiRiT predicts single-cell spatial gene expression using the matched H&E image tiles of human breast cancer and whole mouse pup, evaluated by Xenium (10x Genomics) datasets. Importantly, SPiRiT incorporates rigorous strategies to ensure reproducibility and robustness of predictions and provides trustworthy interpretation through attention-based model explainability. SPiRiT model interpretation revealed the areas, and attention details it uses to predict gene expressions like marker genes in invasive cancer cells. In an apple-to-apple comparison with ST-Net, SPiRiT improved the predictive accuracy by 40%. These gene predictions and expression levels were highly consistent with the tumor region annotation. In summary, SPiRiT highlights the feasibility to infer spatial single-cell gene expression using tissue morphology in multiple-species.

## Introduction

Determining how cells and their states are arranged in space is crucial for grasping how tissue structure relates to its function^1^. The spatial arrangement of cells is also vital in developmental processes and organogenesis in multicellular life forms^2^. Single-cell sequencing necessitates the separation of cells from their native tissues and thus loses important spatial information^3^. The clear advantage of spatial omics technology is that it reveals the spatial organization and potential interactions of cell subpopulations within their native habitats, at a pixel scale ranging from sub-micrometers to 100s of micrometers (µm)^4–14^.

State-of-the-art *in silico* spatial transcriptomics methods predict gene expression across histological images of tissue stained with hematoxylin and eosin (H&E) to characterize cellular heterogeneity at <u>multi</u>-cell resolution (~100 μm). Methods such as ST-Net^15^, HisToGene^16^, DeepSpaCE^17^, Hist2ST^18^, TESLA^19^, BRST-Net^20^, and THItoGene^21^ feature deep learning predictive models trained using H&E images paired with spatial transcriptomics data using the Visium^12^ platform (10x Genomics) to predict expression of hundreds of genes. However, due to Visium’s 55-μm spot size for detecting gene transcripts and the 100□µm center-to-center distance between spots, each spot frequently encompasses several cells. This can hinder the ability to discern intricate tissue architecture and to analyze cell communication patterns, e.g., pinpointing interactions between ligands and receptors. Thus, to leverage the vast amount of commonly available histopathology images for cellular heterogeneity analysis, there is an urgent need for *in silico* single-cell spatial transcriptomic methods. The advent of sub-cellular spatial transcriptomic technologies, e.g., imaging-based RNA hybridization methods (e.g., Xenium)^4–7^ and spatial barcoding-based RNA sequencing methods (Seq-Scope)^9, 10^, offers groundbreaking potential to decode the cellular heterogeneity of tissues including formalin-fixed paraffin-embedded (FFPE)-treated tissue sections.

Among deep learning models, vision transformer (ViT) models have been demonstrated to capture long-range spatial relationships with more robust prediction power for image classification tasks than regular convolutional neural network (CNN) models and also offer better model interpretability^22–24^. Here we leveraged Xenium (10x Genomics) datasets to train ViT models to predict single-cell resolution spatial gene expression from H&E images across multiple species and multiple mouse organs. Our ViT framework, SPiRiT, i.e., Spatial Omics Prediction and Reproducibility Integrated Transformer, maps histological signatures to spatial single-cell transcriptomic signatures.

By integrating cross-validation and model interpretation^25^ for explainable deep learning during hyper-parameter-tuning, SPiRiT accurately predicts single-cell spatial gene expression from histopathological images of human breast cancer and mouse whole pup tissues, evaluated using 10x Genomics Xenium datasets. Our approach demonstrated a 40% improvement in predictive accuracy over the ST-Net CNN algorithm for the ST-Net human breast cancer dataset. These results highlight the capacity of SPiRiT for spatial single-cell gene expression inference across multiple species and organs. Importantly, incorporating explainable deep learning models via model interpretation and vision transformer is expected to contribute as a general-purpose framework for spatial transcriptomics.

## Results

### SPiRiT overview

We developed the SPiRiT framwork using training and testing datasets from single-cell resolution spatial gene expression data processed from the Xenium in Situ platform (10X Genomics) and post-Xenium paired H&E images of the *same* tissue section^6^. The fact that Xenium-assayed tissue sections still retain their histological structure *and thus can be H&E-stained and imaged* affords the opportunity to both train the model and assess its predictive performance, as we can apply it to the post-Xenium H&E image and compare its output to the Xenium data. To enable the model to be effectively applied to related tasks without training from scratch, we used the machine learning approach of transfer learning. This approach allows reuse of a pre-trained model so it can be applied to a new problem, leveraging previously learned knowledge trained on a broad dataset for better generalization on different tasks. The overview of SPiRiT is displayed in **Figure 1**.

**Figure 1.**
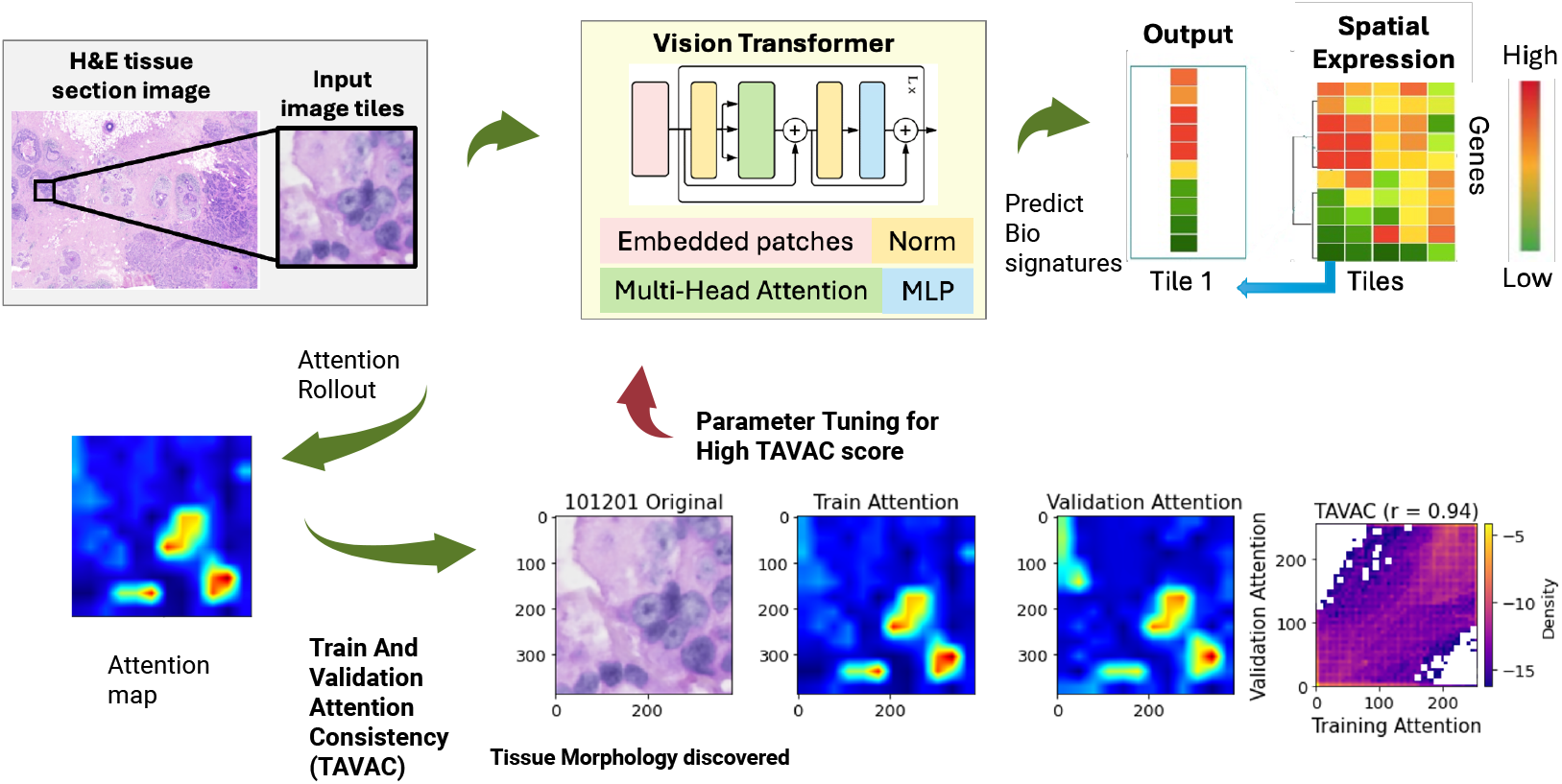
SPiRiT workflow. spot images are extracted from the whole H&E image and used as the input for the Vision transformer models. The Vision transformer will use the spot image to predict the corresponding bio-signatures for the corresponding spot. The signature can be expression levels, pathway scores, etc. TAVAC score is calculated to guide the model selection so that consistent High Attention Region (HAR) is used by ViT no matter the model has seen the data sample during training or not.

SPiRiT uses Xenium resolution tile level histological images as input to predict biological signatures of interest. One challenge in using deep learning models to interpret biomedical images is their inconsistency, as performance can vary significantly based on the data they are trained on^26, 27^. A significant advantage of ViT models is the model’s interpretability afforded by the *self-attention mechanism*, which highlights portions of an image that are critical for prediction (*attention maps*)^28^. However, there is no way to determine *a priori*, based solely on prediction performance, whether the attention maps are trustworthy. SPiRiT is created specifically to handle this problem. SPiRiT is a ViT model that uses consistent image patterns for prediction enhanced by Training and Validation Attention Consistency (TAVAC)^25^. Briefly, TAVAC cross-validates attention maps using the same image in training and testing folds of the model and compares them using a Pearson correlation coefficient. The TAVAC score serves as an independent training metric to ensure that the ViT consistently uses the same image patterns for prediction, regardless of whether the model was trained on the image. Thus, the high-attention regions (HARs) can be used to identify the meaningful biological morphology linked to the biological signature of interest (expression, pathway enrichment, cell types, etc.). In this way, SPiRiT emphasizes the robustness and consistency of model interpretation alongside prediction performance.

### SPiRiT on Xenium breast cancer data identifies the correct cells while predicting cell type marker genes

To test the accuracy and reliability of SPiRiT for predicting human cancer gene biomarkers, we applied SPiRiT to two replicates from the breast cancer Xenium dataset^6^. We used replicate 1 (Rep 1) for training and replicate 2 (Rep 2) for independent validation. After pre-processing and filtering, there were 166,453 cells with corresponding H&E images. To test the ability of SPiRiT to accurately and reliably classify gene expression across Rep 1 and Rep 1, we derived the area under the curve (AUC) from the receiver operating characteristic (ROC) model. We showed the top 5 genes with highest accuracy across Rep 1 (**Figure 2A-B**) and Rep 2 (**Figure 2C-D**), and as expected, with each incremental cut-off filtering less informative predictors (**Figure 2A, C**), the model’s discriminative ability (i.e., AUC) increased.

**Figure 2.**
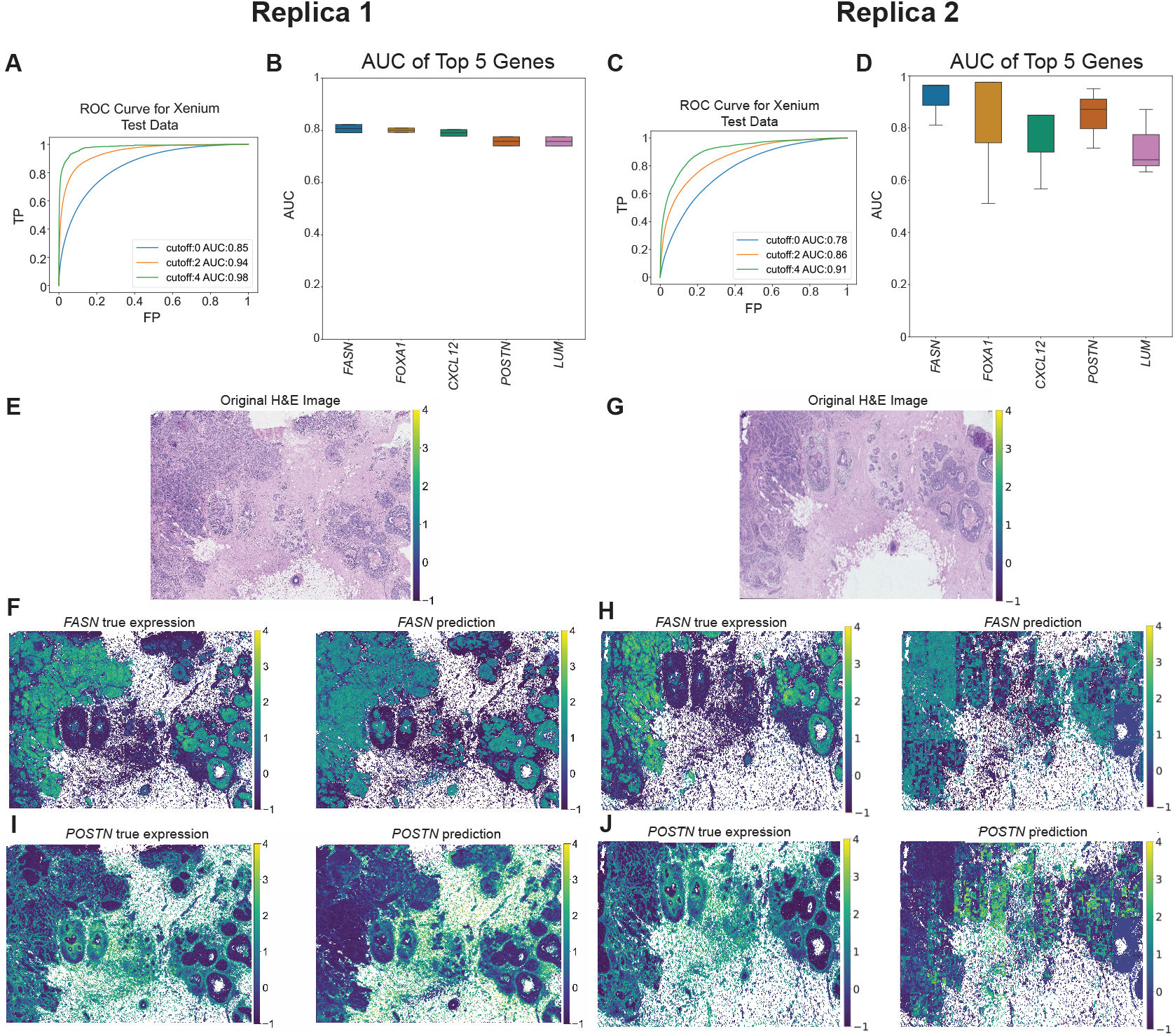
SPiRiT predicts human breast cancer single-cell resolution spatial gene expression from H&E images. (**A**) The ROC curve and AUC value of the ViT model across all genes evaluated using the data from test partition (Replica 1). The plot is generated based on Log(x+1) transformed expression values thus cutoff 0,2,4 corresponds to 1,exp(2), exp(4) transcripts in the cell (**B**) Top 5 genes with highest AUC from Replica 1. **(C)** The ROC curve and AUC value of the ViT model across all genes evaluated using the data from test partition (Replica 2). (**D**) Top 5 genes with highest AUC from Replica 2. (**E**) Original H&E image for Replica 1. (**F**) Heatmap visualization on true expression from Xenium and SPiRiT prediction of *FASN* from Replica 1. (**G**) Original H&E image for Replica 2. (**H**) Heatmap visualization on true expression from Xenium and SPiRiT prediction of *FASN* from Replica 2. (**I**) Heatmap visualization on true expression from Xenium and SPiRiT prediction of *POSTN* from Replica 1. (**J)** Heatmap visualization on true expression from Xenium and SPiRiT prediction of *POSTN* from Replica 2.

SPiRiT predicted genes included previous targets of breast cancer such as fatty acid synthase (*FASN*)^29^ and periostin (*POSTN*)^30^. *FASN* overexpression and hyperactivity in cancers are believed to be obligatory metabolic adaptations, directly selected for their ability to confer growth and survival advantages in the challenging microenvironment of tumors creating a FASN-related lipogenic phenotype necessary for tumorigenesis and metastasis^29^, while POSTN is involved in extracellular matrix remodeling and its overexpression supports epithelial-mesenchymal transition (EMT), a hallmark of metastatic cancers^30^. Other SPiRiT-predicted genes (e.g., FOXA1^31^, CXCL12^32^ and LUM^33^) have also been linked to breast cancer. Moreover, the anti-cancer effects of FASN inhibitors in breast cancer have been extensively documented^34^. As such, we mapped *FASN* and *POSTN* expression back to the H&E slides (**Figure 2E-F**), revealing high SPiRiT predictive accuracy for Rep 1 (**Figure 2G-H**) and Rep 2 (**Figure 2I-J**). SPiRiT’s robust performance occurs despite the inherent section-to-section gene expression variability present in the Xenium dataset^6^, suggesting that SPiRiT can overcome biological noise and experimental variation when using *in situ* gene expression datasets.

Additionally, we examined the root square mean square error (RMSE) to estimate how well SPiRiT performs in terms of prediction accuracy (**Figure S1**). A detailed examination of the top two genes with the lowest RMSE values indicated high degree of zero-inflation in the expression values^35^. Thus, we opted not to use the RMSE for model evaluation for this study’s purposes.

In order to demonstrate the interpretation power of SPiRiT, we further trained SPiRiT to predict a single gene expression. We fine-tuned the ViT with 10,000 randomly selected samples and split the whole data set into two partitions for TAVAC calculation. We picked the marker genes (**Figure 3A**) provided Janesick et al.^6^, namely, *FASN* (invasive cancer cells), *POSTN* (stromal cells), and *IL7R* (lymphocytes). We compared the training attention and validation attention together with the cell type annotations. We found that the HAR, highlighted in red, overlaps with the corresponding marker gene cell type consistently. For instance, in the *FASN* prediction model, the HAR consistently overlaps with the cells that are annotated as the invasive cancer cells (**Figure 3B**, red in column 2). We obtained comparable results for the *POSTN* (**Figure 3C**). and *IL7R* (**Figure S2)**. As such, SPiRiT can be used to link biological signatures with tissue morphology and quantitively measure the confidence level of the association in human tissue.

**Figure 3.**
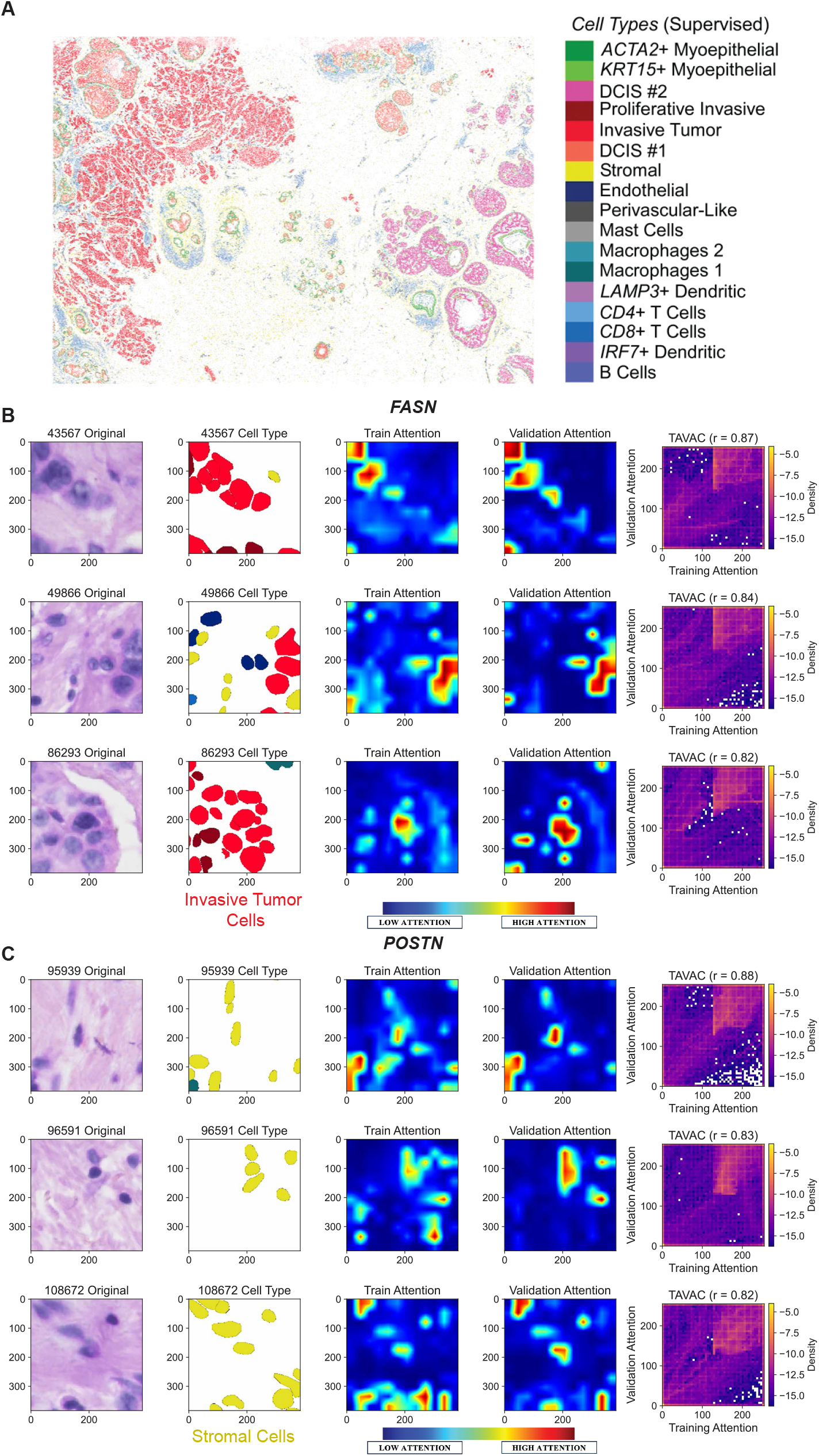
High Attention Region (HAR) can be used to identify corresponding cell types. (**A**) Cell type annotation for Xenium breast cancer data. (**B-C**) H&E image and cell type annotation using SPiRiT for *FASN* and *POSTN*, respectively. Each row shows one data point. The original H&E tile image is shown in the first column. In the second column, the corresponding cell type annotation is displayed. The Training Attention and Validation Attention are shown respectively on column 3 and 4. The HARs (red regions) when the ViT model predicting the gene expression for *FASN* and *POSTN*, overlap with the corresponding invasive tumor cells (red cells in **Figure 3A**) and stromal cells (yellow cells in **Figure 3A**), respectively. TAVAC score calculation is visualized as scatter plot between Training Attention and Validation Attention on column 5.

### SPiRiT on Xenium mouse data predict whole pup in situ gene expression

To evaluate the generalization of our framework across multiple species, we further trained SPiRiT model using the whole mouse pup *in situ* gene expression using the Xenium mouse tissue atlassing panel. The SPiRiT model (input: H&E image, output: gene expression at ~4 µm) positively predicted expression of all Xenium assayed genes (n = 379) in the test partition (**Figure 4A**). The ROC curve supports the robustness of the SPiRiT model across all genes evaluated using the data from test partition (**Figure 4B)**. The top 20 predicted genes also showed high model’s accuracy (AUC ranging between 0.90 to 1.0; **Figure 4C**) and test-retest correlation (**Figure 4D**). The gene with the highest prediction vs. truth correlation, *Ttn*, a smooth muscle gene marker, shows a high correlation (r = 0.7) between the Xenium spatial gene expression measurement and SPiRiT model prediction across the spot of the whole mouse pup body (**Figure 4E**). The *Ttn* gene provides instructions for making a large protein called Titin, which plays an important role in skeletal and cardiac (heart) muscle function^36^. In agreement with this, high expression of *Ttn* was observed and accurately predicted using SPiRiT in regions of skeletal muscle (**Figure 4F**). *Pygm*, encoding muscle glycogen phosphorylase, also revealed similar patterns between true expression and prediction heatmaps (**Figure 4G)**. These results demonstrate that SPiRiT offers high predictive accuracy and reliability, making it suitable for predicting *in situ* gene expression of mouse tissue.

**Figure 4.**
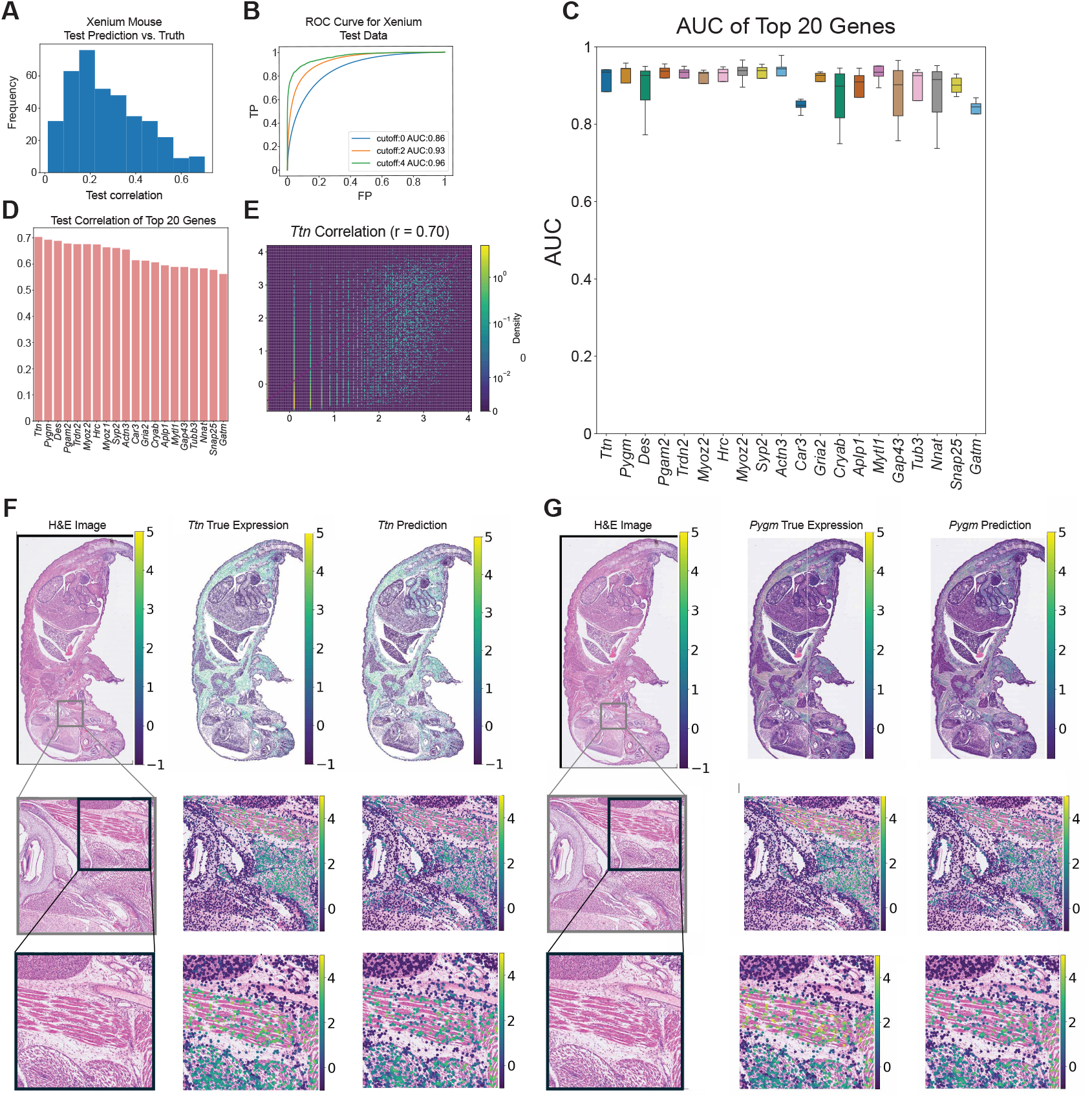
SPiRiT predicts mouse single-cell resolution spatial gene expression from H&E images. (**A**) Performance of the Xenium-based ViT model is demonstrated by the histogram of correlation coefficients of prediction vs true expression for each gene on mouse Xenium test data. Training and test data are Xenium In Situ data of the expression of 379 genes and post-Xenium paired H&E images, aligned by Fiji BigWarp (v9.0.0). (**B**) The ROC curve and AUC value support the robustness of the ViT model across all genes evaluated. (**C-D**) Top 20 genes with the highest predicting performance measured by classification and correlation between measured and predicted gene expression levels, respectively. (**E**) Scatter plot with true expression vs predicted expression values on Xenium test data for gene *Ttn*. **(F**) H&E images, the *Ttn* gene expression via Xenium measurements (middle panels) and via ViT model prediction (right panels) of mouse whole body section and zoomed-in regions. **(G**) H&E images, the *Pygm* gene expression via Xenium measurements (middle panels) and via ViT model prediction (right panels) of mouse whole body section and zoomed-in regions.

### SPiRiT trained on spatial multi-cell transcriptome data outperform CNN model

We further validated the predictability and generalizability of SPiRiT using an independent breast cancer spatial transcriptomic data and matched H&E images from the same tissue section (ST-Net dataset)^37^. The ST-Net dataset contains 30,612 spatially mapped gene expression profiles generated from tumor specimens from 23 persons with breast cancer. To facilitate a direct comparison with ST-Net, the SPiRiT model uses H&E-stained breast cancer sections as the input, while the output is the *log*(*x* + 1) transformed gene expression from each corresponding pixel (spot) of the image. As in^37^, only the 250 genes with the highest mean expression levels were included for the prediction. Leave-one-patient-out cross validation was performed where we used 22 patients’ data for training and the left-out one patient data for testing. We measured the correlation between prediction and real expression values on test patients. The metric ST-Net used for performance evaluation is the number of genes predicted: the number of genes with positive prediction vs real correlation for more than 20 out of 23 patients. The results from the ST-Net dataset show that the pretrained SPiRiT model had the best performance and surpassed ST-Net by 40% (**Figure 5A**). Note that even the *de novo* SPiRiT had better performance than ST-Net pretrained models. This case-series analysis shows that SPiRiT has stronger prediction power than state-of-the-art convolutional neural networks.

**Figure 5.**
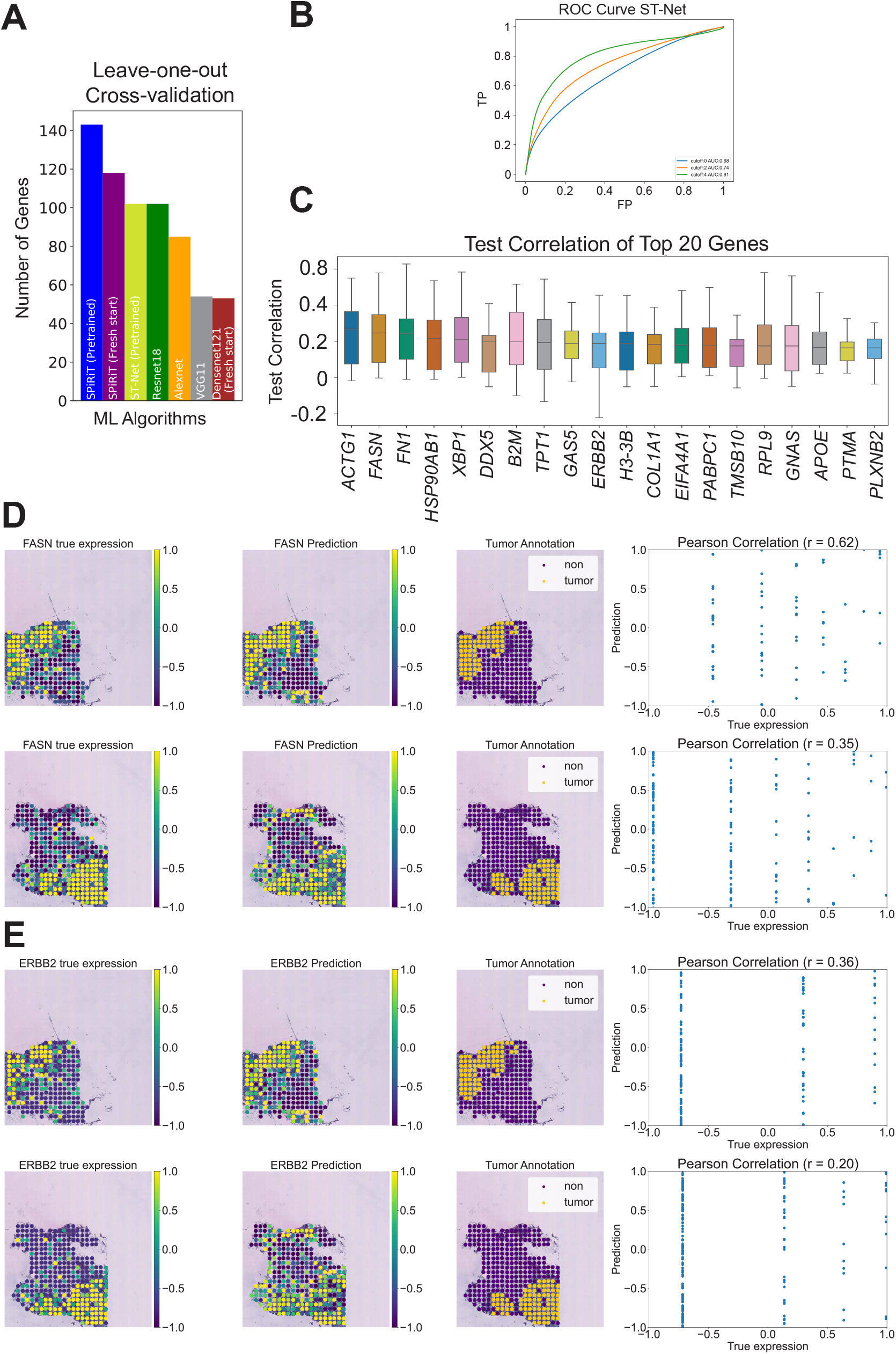
SPiRiT predicts human multi-cell resolution spatial gene expression from H&E image. (**A**) The number of genes exhibit positive correlation between predicted and measured gene expression across deep learning architectures. SPiRiT (ViT) outperforms other methods: ST-Net, Resnet18, Alexnet, VGG11 and Densenet121. The best performance is achieved for SPiRiT pretrained with ImageNet dataset (blue bar). The fresh start SPiRiT achieves second best performance. (**B**) The ROC curve and AUC value support the robustness of the ViT model across all genes evaluated. (**C-D**) Top 20 genes with the highest predicting performance measured by correlation between measured and predicted gene expression levels. **(D-E)** Cancer biomarker *FASN* and *ERBB2* expression map vs prediction and tumor annotation of two tissue sections, respectively. The prediction and expression levels are highly consistent with the tumor region annotation (yellow region).

To test whether SPiRiT can accurately and reliably predict cancer spatial gene expression in the ST-Net dataset, we performed ROC models and test-retest correlation analyses. Indeed, as shown with the Xenium human breast cancer (**Figures 2A-D**) and whole mouse (**Figures 2B-C**) spatial gene expression datasets, SPiRiT showed high predictive accuracy when applied to the ST-Net dataset (**Figure 5B-C**). The top 10 predicted genes with highest median correlations of all test patients were all cancer biomarkers. *FASN*, as described above, has been extensively shown to be a therapeutic target in breast cancer^34^. It was accurately and reliably predicted by Janesick et al.^6^, our SPiRiT model using two separate replicas from the Xenium human breast cancer dataset (**Figures 2E-J**), the original ST-Net model^37^ and our SPiRiT application to ST-Net dataset (**Figure 5D**). This consistent detection across multiple datasets suggests a significant and reproducible role of FASN in the pathogenesis of breast cancer, and SPiRiT’s ability to discover and identify biologically relevant cancer biomarkers.

Additionally, SPiRiT predicted highly relevant breast cancer genes among its top 10 hits. *EBBR2* (**Figure 5E**), also known as *HER2*, overexpression is commonly found in individuals with breast cancer, and its mutations are linked with the more aggressive, chemo-resistant and recurrent HER2-positive breast cancer subtype^38^. *XBP1* (**Figure 5F**) has been shown to promote triple-negative breast cancer^39^ and induce EMT^40^, while its inhibition has been shown to suppresses breast cancer^41^. Other SPiRiT-identified genes, e.g., *FN1* (**Figure S3A**) and *COL1A1* (**Figure S3B)** have also been reported to strongly breast cancer^42–45^. *FASN, GNAS, ACTG1* and *DDX5* are also among the top 5 well predicted genes by ST-Net^37^. *GNAS* (**Figure S4A**) mutations have been found in various types of tumors, including pituitary, thyroid, and pancreatic^46^. *ACTG1* (**Figure S4B**) has been implicated in tumorigenesis and metastasis in various cancers^47^. *DDX5* (**Figure S4C**), has been implicated in various cancers, including breast and lung cancer^48^. Importantly, SPiRiT’s predictive correlation remained consistent across different technical replicates, suggesting high reproducibility overcoming biological and experimental variation. Collectively, these results suggest that SPiRiT is able to identify biologically meaningful and relevant cancer biomarkers with higher accuracy and reproducibility than the original ST-Net model.

## Discussion

The application of deep learning to spatial omics has ushered in a new era of high-resolution tissue profiling, enabling the deconvolution of cellular heterogeneity and spatial gene expression at unprecedented scale and accuracy. Our current work leverages the potential of ViT and TAVAC to decode the complex interplay between gene expression and cellular morphology through a newly developed model named ‘SPiRiT’. Here, we introduce SPiRiT, a vision transformer (ViT)-based framework that leverages histological image features to predict spatial single-cell transcriptomic signatures. By systematically evaluating SPiRiT on human breast cancer and whole mouse pup datasets, including two independent Xenium breast cancer replicates (**Figure 2**) and a whole mouse pup dataset^6^ (**Figures 4**), we demonstrate robust and reproducible prediction of disease-relevant and non-disease gene biomarkers.

SPiRiT’s performance, notably a 40% improvement in predictive accuracy over ST-Net^37^ establishes a new benchmark for *in silico* spatial transcriptomics. The model’s predictions were highly concordant with expert tumor region annotations, underscoring its potential utility in digital pathology and biomarker discovery. Importantly, our adoption of TAVAC for hyperparameter tuning and model interpretability addresses a critical challenge in the deployment of deep learning for clinical applications: the need for transparent, reproducible, and reliable attention mechanisms. By enforcing a stringent TAVAC threshold, we ensured that only models with consistent and interpretable attention patterns were advanced, a consideration of relevance in medical diagnostics where reproducibility is paramount.

Different from other computational approaches such as xFuse^49^ and iStar^50^, which require Visium data as input, SPiRiT directly utilizes H&E-stained histological images, broadening its applicability and facilitating integration with routine clinical workflows. This flexibility positions SPiRiT as a promising tool for personalized medicine, enabling precise mapping of transcriptomic signatures to imaging signatures to inform individualized therapeutic strategies. The capacity to delineate cancer biomarkers within heterogeneous tumor microenvironments may ultimately guide precision oncology interventions and improve patient outcomes.

The translational potential of SPiRiT is further highlighted by its successful application across species and tissue types, including whole-body mouse sections, thus extending the utility of deep learning models beyond human cancer to broader biological contexts. This cross-species generalizability aligns with recent advances such as the SCHAF model^51^, but SPiRiT’s demonstrated efficacy in both human and mouse datasets represents a significant step forward.

Nevertheless, our approach has certain limitations. While SPiRiT generalizes well across the datasets tested, further validation across additional tissue types, disease contexts, and imaging modalities is warranted. Additionally, the computational resources required for training vision transformers may present a barrier for some laboratories, though ongoing advances in model efficiency are likely to mitigate this concern. Looking ahead, SPiRiT opens new avenues for research and clinical application. Its ability to map transcriptomic signatures to tissue morphology with high precision suggests potential for guiding personalized treatment strategies and supporting rapid, automated histopathological diagnostics. Future work will focus on extending SPiRiT to additional species and organs, integrating multi-omics data, and exploring its utility in prospective clinical studies. In summary, SPiRiT exemplifies the synergy of computational innovation and biological insight, providing a robust and interpretable framework for spatial transcriptomics that is poised to advance both basic research and precision medicine.

## Methods

### Vision Transformer Model architecture

ViT^23^, a pioneering machine learning model developed by Google Brain, marks a notable departure from the conventional application of convolutional networks in visual recognition tasks. Rather, it adopts the Transformer model, traditionally utilized in natural language processing, and applies it to image processing. In this architecture, images are deconstructed into sequences of patches, akin to the treatment of text as sequences of words or characters within the domain of language. Each image patch is linearly embedded and then interpreted as an input token, with the addition of position embeddings to preserve spatial information. Subsequently, the model is trained end-to-end through the utilization of self-attention and feed-forward networks, which empowers the architecture with the capacity to model intricate, long-range interactions amongst patches within an image. This innovative approach has yielded substantial advancements in numerous image classification tasks, hence attesting to the robustness and versatility of Transformer models beyond the scope of language comprehension.

### Pretrained Vision Transformer Model architecture

The ViT model, specifically the ViT-L/32 configuration, trained on the ImageNet-1k dataset, is a comprehensive and layered architecture intended for image classification tasks. It adopts the architecture principles of Transformer models, typically used in the realm of natural language processing, and adapts it to handle imaging data.

In the ViT-L/32 configuration, ‘L’ stands for ‘Large’, indicating a larger model variant. It comprises 24 transformer layers, each embedding 1024 nodes. An image is divided into a grid of non-overlapping patches, where each patch size is 32×32 pixels. The flattened patch is then passed through a linear projection to generate a 1024-dimensional embedding. These embeddings are processed as input tokens for the Transformer model. The model also introduces a special learnable [CLS] token at the beginning of the sequence of patch embeddings, which is later used for classification. Position embeddings are added to these image patch embeddings to incorporate spatial information, which is crucial in image understanding tasks. Each transformer layer in the architecture consists of a multi-head self-attention mechanism and a two-layer feed-forward neural network, facilitating the model’s ability to learn complex, long-range interactions among the image patches. The final output of the model is obtained by applying a Multi-Layer Perceptron (MLP) to the [CLS] token’s representation.

The MLP consists of two fully connected (FC) layers with 512 hidden nodes each. The FC layer has nonlinear activation function: Rectified Linear Unit (ReLU). Then the output is followed by a dropout layer (drop rate = 0.5). The final output is softmax layer (classification output) or a 3rd FC layer (regression output) depends on the output data type. In the leave one patient out cross validation experiment, FC layer is used since the output is the log gene expression of 250 genes.

### Leave-one-patient-out Cross Validation on ST-Net data

In our research, the Vision Transformer models were systematically trained with the objective of predicting the expression levels of the 250 genes demonstrating the highest mean expression values. An examination of these genes revealed that a considerable number exhibited extremely low expression levels as shown in ST-Net paper^37^. Consequently, this led to the predicament of a low signal-to-noise ratio, which significantly complicated the prediction process.

To ascertain the generalizability of our predictions to unseen samples, we adopted a rigorous leave-one-out cross-validation method. This entailed iterative training on datasets from 22 patients while predicting the gene expression of the remaining patient. For each cross-validation fold, the model was initialized utilizing pre-trained ImageNet weights, and all weights were trained employing stochastic gradient descent, with a learning rate of 0.0001 for pretrained ViT and 0.000001 for customized ViT. Pretrained ViT is trained for 25 epochs while ViT with fresh starts takes 1500 epochs. The model operated with a batch size of 128 (pretrained) and 256 (fresh start).

During the training phase, the dataset was expanded by randomly rotating each image by 0, 90, 180^°^, and applying a random flip vertically or horizontally 50% of the time. Center crop image augmentation is also introduced during training.

### Attention interpretation

We used attention rollout^28^ to generate the attention heatmap given an input image. Attention rollout is a technique that expands on the concept of attention in transformer models, utilized to provide interpretability to these models, which are typically characterized as “black boxes”. Given the nature of attention in transformer models where each layer has its own attention distribution, simple aggregation methods may not sufficiently capture the multi-layer, multi-head attention complexity. To overcome this limitation, attention rollout computes an approximation of the aggregated attention by rolling out the attention process over multiple layers in a sequential manner. It calculates the contribution of every input token to every output token by traversing the attention mechanism from the last layer to the first. This process generates a singular attention matrix that embodies the aggregated influence of all attention heads across all layers, thus yielding an interpretable summary of the original model’s decision-making process. As such, attention rollout contributes significantly to our understanding of how deep transformer models operate and make predictions, thereby advancing the field’s interpretability efforts. A filter on attention maps is also introduced where only the top 10 percents of high attention values are used. The original attention rollout paper suggests taking the average attention across the attention heads, but it seems taking maximum value instead generates better results. The result reported in this paper uses max function to aggregate the values from attention heads.

### Train And Validation Attention Consistency (TAVAC): ranking system to quantify how trustworthy your Vision Transformer model is

We propose the Train and Validation Attention Consistency (TAVAC) score. The TAVAC score is calculated by the following process: Using a similar idea as K fold cross validation, we divided the given data set D into two folds. In the first stage, we use one-fold of data as a train and the other as a validation set. We subsequently trained the model on the training dataset and evaluated it on the validation dataset. In the second stage, we switch the roles of our two subsets, making the training dataset in stage 1 validation and the validation dataset in stage 1 training. Again, we trained the model on the training dataset and evaluated its performance on the validation dataset. Finally, we examined the model’s prediction power and consistency by comparing corresponding attention results for images in the stage 1 training against stage 2 validation. In each stage, we generate the attention map for each image using Attention Rollout ^28^. If an image is used as training data, the generated attention map will be defined as Training Attention. If the image is used as validation data, the attention map will be defined as Validation Attention. The consistency between Train and Validation Attention is defined as Train and Validation Attention Consistency (TAVAC) score. In this study, we used Pearson Correlation to quantify the consistency. It can be extended to other consistency scores in the future.

### Using TAVAC to direct SPIRIT training

Here, we introduce an innovative approach to the training of the SPiRiT deep learning model, prioritizing not just the performance metrics typically observed, such as accuracy and F-1 score, but also the consistency of features throughout the training process. To achieve this balance, we employ the TAVAC score^25^ as a criterion for hyper-parameter tuning, ensuring that the patterns utilized for prediction by the ViT are reliable and consistent. This leads to a trade-off scenario where a high TAVAC score must be carefully weighed against the model’s predictive performance. Our current strategy is stringent: we dismiss any parameter settings that result in a median TAVAC score below 0.7. From the remaining models, we then select the one with the highest TAVAC score that still delivers acceptable prediction performance. This methodology underscores the importance of feature stability, aiming to enhance the interpretability of the model’s decision-making process.

### SPiRiT training for Xenium breast cancer data

With Xenium data, cell perimeters are delineated within the images before attributing transcripts to specific cells, mirroring the cell identification phases in single-cell transcriptomics^6^. This process involves identifying nuclei via DAPI staining, then extending these regions up to a 15 μm maximum or until encountering adjacent cell borders. In the analyzed sample, a total of 167,885 cells and 36,944,521 transcripts were identified, with an average of 166 transcripts per cell^6^. We extracted H&E images for each cell and used the images to predict the 313 gene expression values for each cell. Some extracted images contained only black background and thus were be filtered out. After filtering we had 166453 cells. All gene expression was log(x+1) transformed. The whole data set was divided into training validation and test partitions with the ratio of 3:1:1. An initial learning rate of 0.0001 is used with linear decay (0.5) every 50 epochs. Weight decay of 0.0001 is applied for regularization. The model is trained for 1000 epochs with batch size of 64. Mean square error (MSE) is used as the loss function. The loss achieves 0.17 for all three partitions. Pearson correlation between predicted and true expression is used for evaluation.

### SPIRIT training for Xenium mouse data

The paired H&E image and Xenium dataset from the same tissue section contains over 1.3 million cells and 191.4 million high quality transcripts. The 379 genes measure by the Xenium panel were sequenced using the FFPE protocol. For each cell, H&E images were extracted and used to predict the expression values of these 379 genes. Images containing only a black background were filtered out, leaving 1.3 million cells for analysis. A subset of 160,000 cells was randomly selected, yielding good performance. All gene expression data was log(x+1) transformed. The data set was divided into training, validation, and test partitions in a ratio of 3:1:1. An initial learning rate of 0.0001 with linear decay (0.5) every 50 epochs was employed, along with a weight decay of 0.0001 for regularization. The model was trained for 1000 epochs with a batch size of 64, using mean square error (MSE) as the loss function. The loss achieved 0.13 across all three partitions. Pearson correlation between predicted and true expression values was used for evaluation.

### Benchmark datasets

We used two primary datasets for benchmarking: Xenium *in situ* data of human breast cancer samples and mouse tissue samples.

For the human breast cancer data, we utilized Sample 1, Replicate 1 from a previous study^6^. This dataset, referred to as Human Breast Cancer (Sample 1), comprises 167,885 cells and 36,944,521 transcripts, with an average of 166 transcripts per cell. The data was processed to extract H&E images for each cell, filtering out images with only black backgrounds, resulting in 166,453 cells for analysis. Gene expression data was log(x+1) transformed. More details can be found in original study^6^.

For the mouse data, we used Xenium In Situ data generated using the Mouse Tissue Atlassing panel on a one-day-old mouse pup (C57BL/6 strain) obtained from AcePix Biosciences (Xenium whole mouse data). This dataset, referred to as Mouse Pup, included 1,355,849 cells and 191,407,876 high-quality transcripts, with a median of 113 transcripts per cell. The tissue section was prepared according to standard FFPE protocols.

Additionally, we analyzed the ST-Net spatial transcriptomics dataset^37^, which captures the spatial distribution of messenger RNA (mRNA) sequences within tissue sections. This dataset consisted of 23 breast cancer patients, with three microscope images of H&E-stained tissue slides and the corresponding spatial transcriptomics data for each patient. Spatial transcriptomics measures RNA expression in spots with a diameter of 100 μm arranged in a grid with a center-to-center distance of 200 μm. Each spot captures several thousand mRNA sequences, represented as a 26,949-dimensional vector of non-negative integers, indicating the number of times a gene was counted. The dataset included 26,949 distinct mRNA species across all samples. Ethical regulations were complied with for experiments involving human tissue samples, and detailed protocols were followed for RNA-seq and subsequent analysis.

## Acknowledgments

Dr. Sheng Li is supported by the following grants: R35GM133562, U01HG013175, U01CA271830. Dr. Matthew Mahoney is supported by R01GM141309. Drs. Matthew Mahoney and Sheng Li are supported by U54AG079753, as the NIH Common Fund Cellular Senescence Network Consortium JAX-Sen Mouse Tissue Mapping Center (TMC). We would like to thank Drs. Nadia Rosenthal, Ron Korstanje, Paul Robson for scientific discussion and feedback. We would also like to express our sincere gratitude to senior scientific writer Carmen Robinett, whose invaluable contributions significantly enhanced the quality of this manuscript.

## Code and Data Availability

The code is available in github: https://github.com/LabShengLi/SPiRiT.

Xenium human breast cancer data: https://www.10xgenomics.com/products/xenium-in-situ/preview-dataset-human-breast

Xenium whole mouse data: https://www.10xgenomics.com/datasets/mouse-pup-preview-data-xenium-mouse-tissue-atlassing-panel-1-standard

ST-net data is downloaded at: https://data.mendeley.com/datasets/29ntw7sh4r/5

