## Supplementary material for "Inferring single-cell spatial gene expression with tissue morphology via explainable deep learning": Figure S1, Figure S2, Figure S3, Figure S4

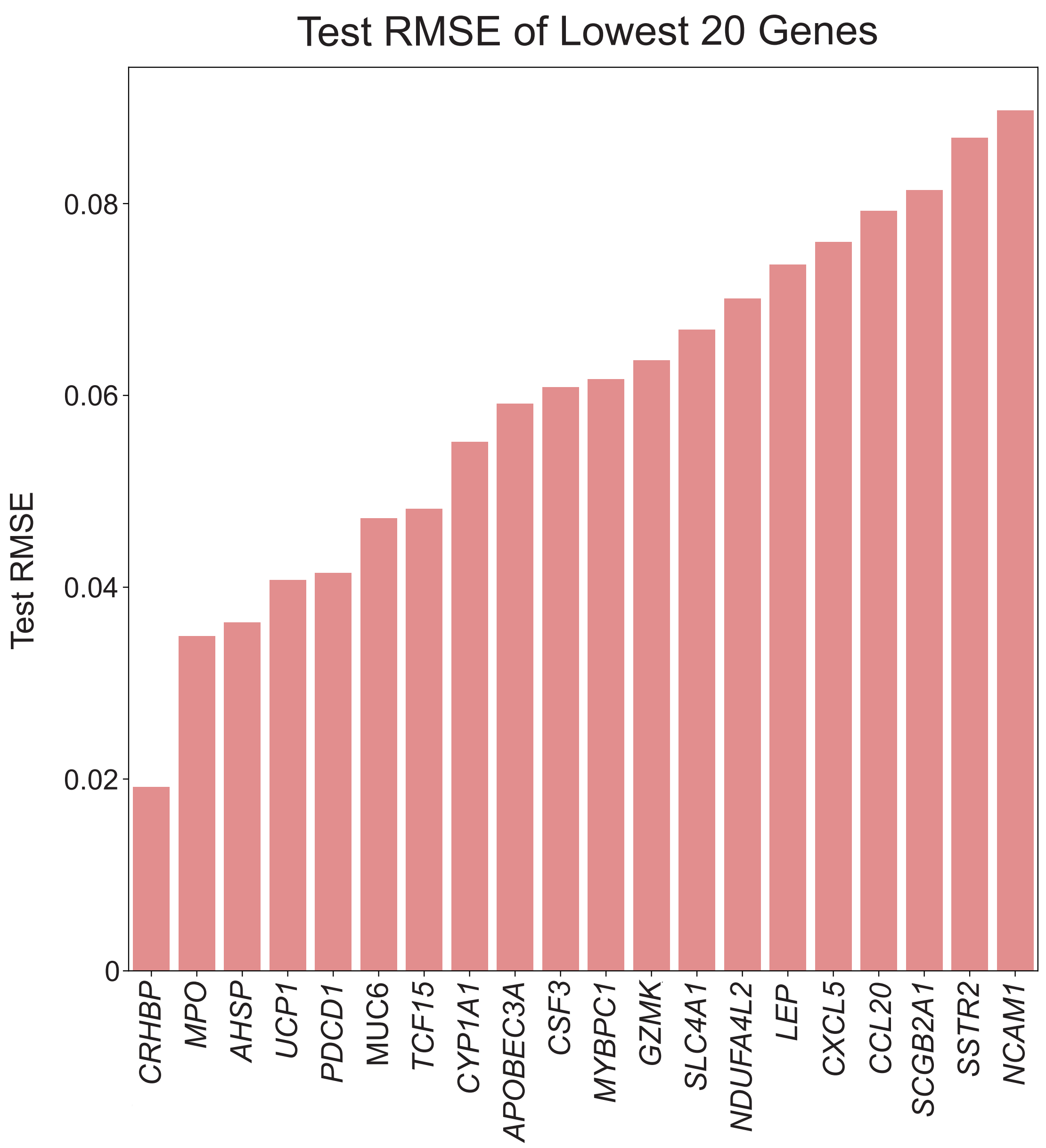


**Figure S1. Top 20 genes with the lowest RMSE between Xenium measured gene expression and SPiRiT predicted gene expression.**

# *IL7R*

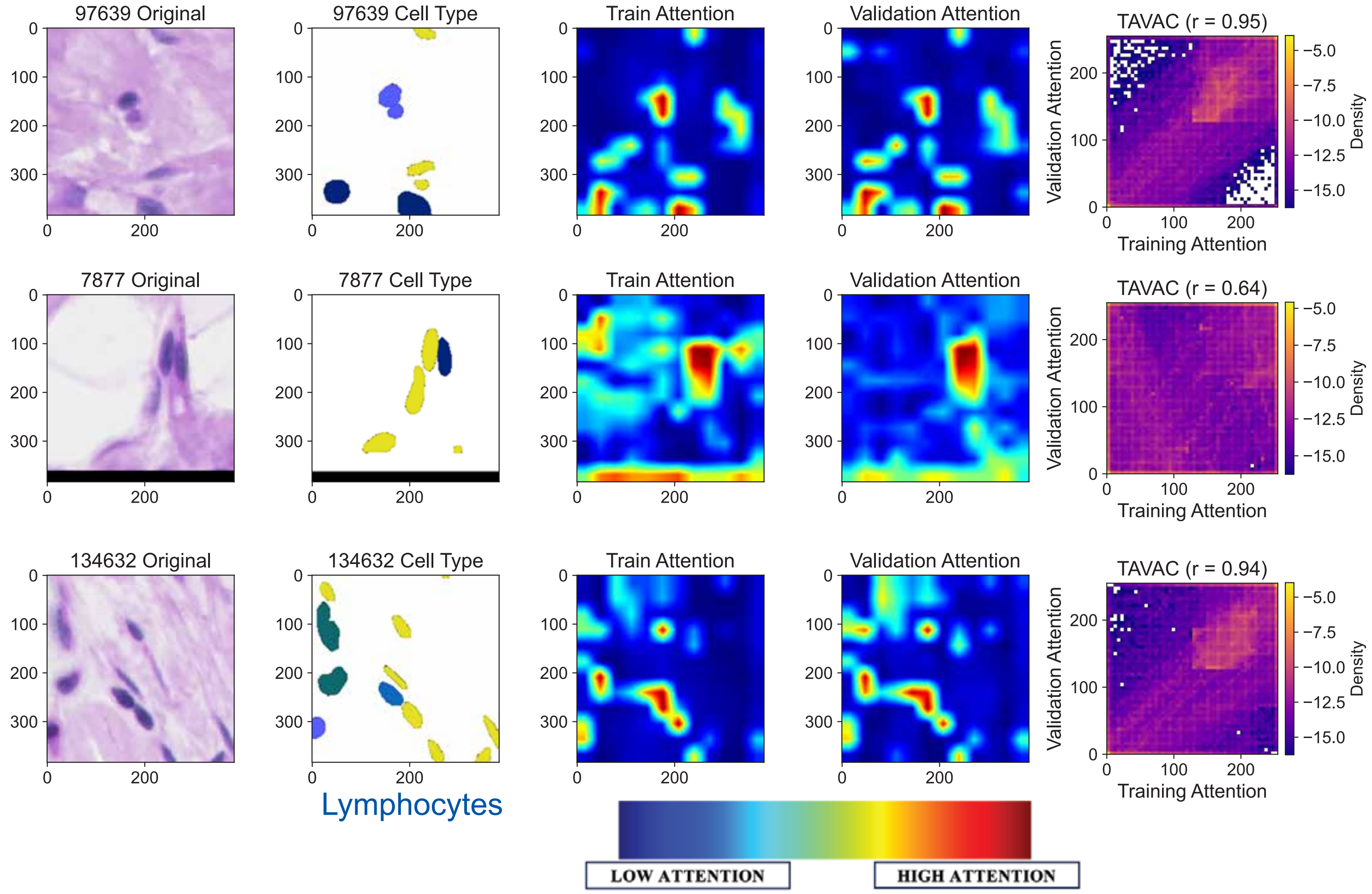


**Figure S2. High Attention Region (HAR) can be used to identify corresponding cell types.** H&E image and cell type annotation using SPiRiT for *IL7R;* Each row shows one data point. The original H&E tile image is shown in the first column. In the second column, the corresponding cell type annotation is displayed. The Training Attention and Validation Attention are shown respectively on column 3 and 4. The HARs (red regions) when the ViT model predicting the gene expression for *IL7R,* overlap with the corresponding invasive tumor cells (blue cells in **Figure 3A**). TAVAC score calculation is visualized as scatter plot between Training Attention and Validation Attention on column 5.

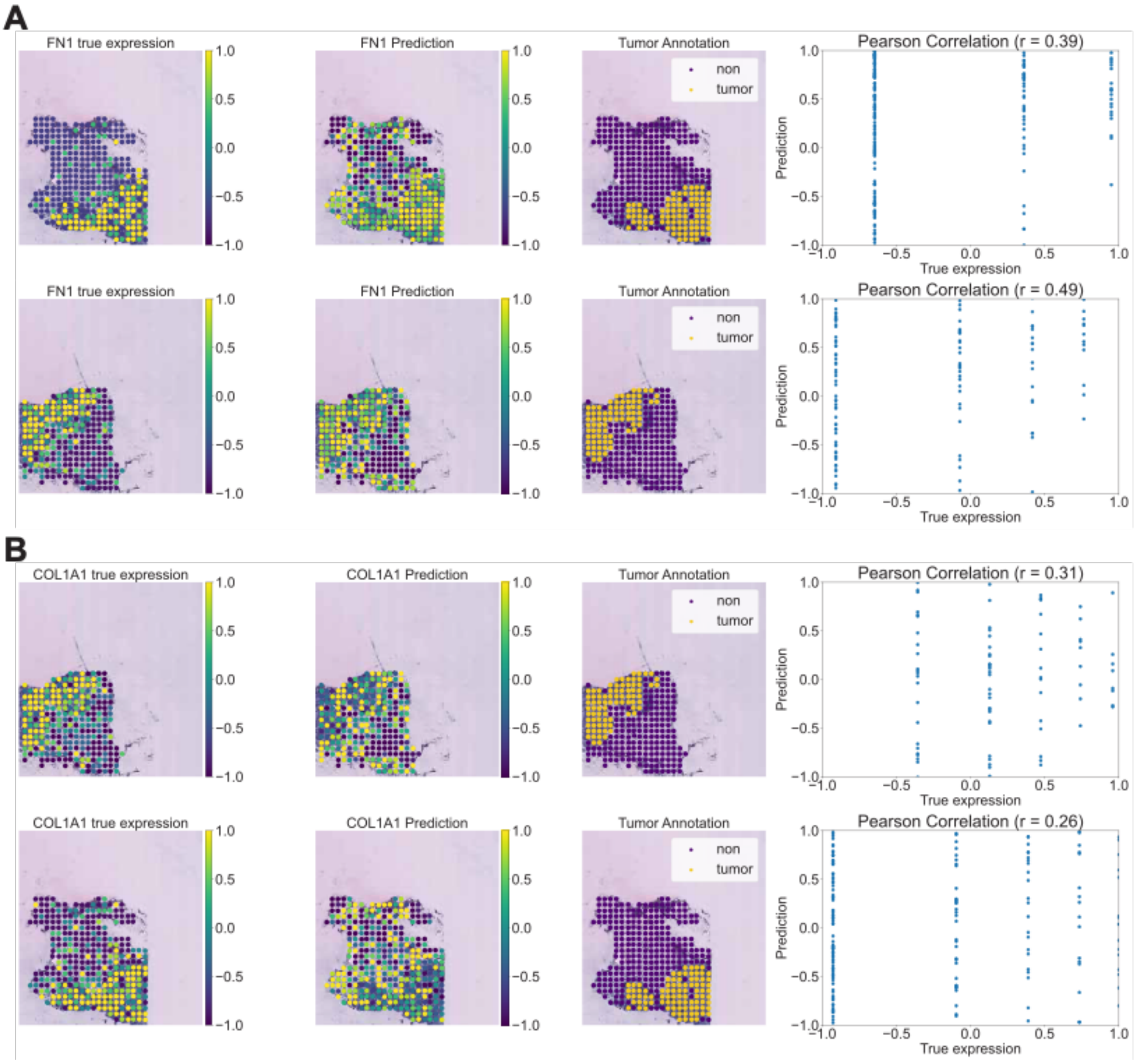


**Figure S3. SPiRiT predicts additional multi-cell resolution spatial gene expression from H&E images. (A-B)** Cancer biomarker *FN1* and *COL1A1*, respectively, expression map vs prediction and tumor annotation of two tissue sections, respectively. The prediction and expression levels are highly consistent with the tumor region annotation (yellow region).

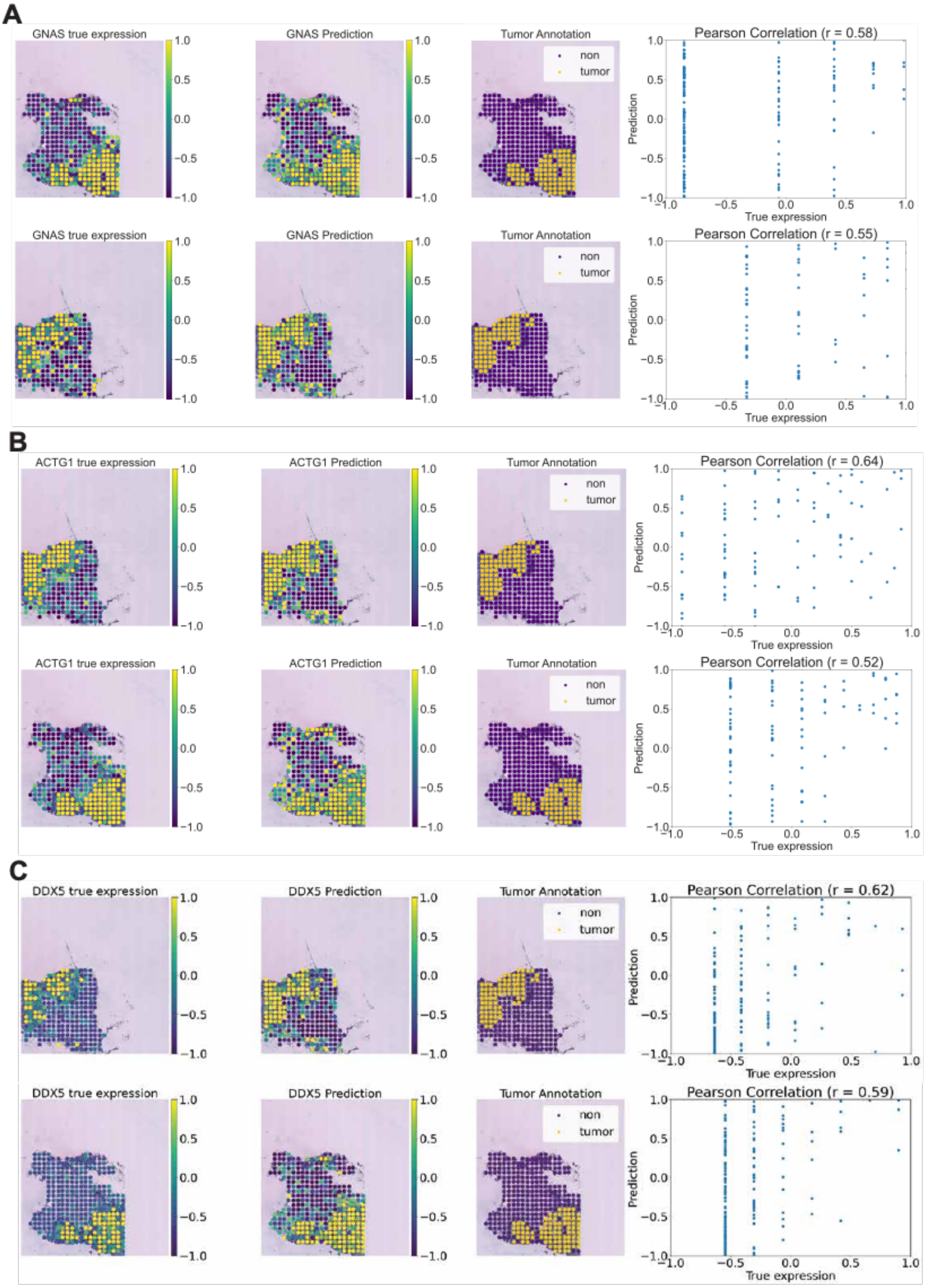


**Figure S4. SPiRiT predicts additional multi-cell resolution spatial gene expression from H&E images. (A-C)** Cancer biomarker *GNAS, ACTG1* and *DDX5,* respectively, expression map vs prediction and tumor annotation of two tissue sections, respectively. The prediction and expression levels are highly consistent with the tumor region annotation (yellow region).
